# Within-patient genetic diversity of SARS-CoV-2

**DOI:** 10.1101/2020.10.12.335919

**Authors:** Jack Kuipers, Aashil A Batavia, Kim Philipp Jablonski, Fritz Bayer, Nico Borgsmüller, Arthur Dondi, Monica-Andreea Drăgan, Pedro Ferreira, Katharina Jahn, Lisa Lamberti, Martin Pirkl, Susana Posada-Céspedes, Ivan Topolsky, Ina Nissen, Natascha Santacroce, Elodie Burcklen, Tobias Schär, Vincenzo Capece, Christiane Beckmann, Olivier Kobel, Christoph Noppen, Maurice Redondo, Sarah Nadeau, Sophie Seidel, Noemie Santamaria de Souza, Christian Beisel, Tanja Stadler, Niko Beerenwinkel

## Abstract

SARS-CoV-2, the virus responsible for the current COVID-19 pandemic, is evolving into different genetic variants by accumulating mutations as it spreads globally. In addition to this diversity of consensus genomes across patients, RNA viruses can also display genetic diversity within individual hosts, and co-existing viral variants may affect disease progression and the success of medical interventions. To systematically examine the intra-patient genetic diversity of SARS-CoV-2, we processed a large cohort of 3939 publicly-available deeply sequenced genomes with specialised bioinformatics software, along with 749 recently sequenced samples from Switzerland. We found that the distribution of diversity across patients and across genomic loci is very unbalanced with a minority of hosts and positions accounting for much of the diversity. For example, the D614G variant in the Spike gene, which is present in the consensus sequences of 67.4% of patients, is also highly diverse within hosts, with 29.7% of the public cohort being affected by this coexistence and exhibiting different variants. We also investigated the impact of several technical and epidemiological parameters on genetic heterogeneity and found that age, which is known to be correlated with poor disease outcomes, is a significant predictor of viral genetic diversity.

**Author Summary:** Since it arose in late 2019, the new coronavirus (SARS-CoV-2) behind the COVID-19 pandemic has mutated and evolved during its global spread. Individual patients may host different versions, or variants, of the virus, hallmarked by different mutations. We examine the diversity of genetic variants coexisting within patients across a cohort of 3939 publicly accessible samples and 749 recently sequenced samples from Switzerland. We find that a small number of patients carry most of the diversity, and that patients with more diversity tend to be older. We also find that most of the diversity is concentrated in certain regions and positions of the virus genome. In particular, we find that a variant reported to increase infectivity is among the most diverse positions. Our study provides a large-scale survey of within-patient diversity of the SARS-CoV-2 genome.

## Introduction

Severe acute respiratory syndrome coronavirus 2 (SARS-CoV-2), the cause of COVID-19, spread globally from its origins in Wuhan, China, toward the end of 2019, resulting in the World Health Organisation declaring COVID-19 a pandemic in March 2020. Initially diagnosed as a pneumonia of unknown origin, huge research efforts have drastically developed our clinical knowledge of COVID-19 concerning its etiology [1], mode of transmission [2], identification of individuals at risk [3] and potential treatment strategies [4].

COVID-19 is a respiratory disease marked by a variety of symptoms including a persistent cough, fatigue and anosmia [5], however, a subpopulation of SARS-CoV-2-infected individuals remains asymptomatic [6]. The large variation in the severity of experienced symptoms, along with the virus incubation time ranging from 5-14 days [7, 8], may have contributed to its rapid propagation. At the end of September 2020, there have been more than 34 million confirmed cases with over a million deaths worldwide [9].

SARS-CoV-2 belongs to the family *Coronaviridae* and is the latest of three zoonotic coronaviruses that have spilled over to infect humans in the past two decades following SARS-CoV in 2003 and MERS-CoV in 2012 [10]. The SARS-CoV-2 single-stranded RNA genome consists of at least 13 open reading frames (ORFs) spanning 29,903 nucleotides [11]. ORF1ab codes for a polyprotein from which the non-structural proteins originate, including the RNA-dependent RNA polymerase; this is then followed by the Spike (S) gene, the Envelope (E) gene, the Matrix (M) gene, the Nucleocapsid (N) gene and a host of accessory genes [12]. Much like the related SARS-CoV, the Spike protein present on the surface of the SARS-CoV-2 virion is central for its access to the target host cell via binding of the hACE2 receptor, S protein priming and finally fusion of the viral and target cell membranes [12–14]. Zeigler et al. identified co-expression of hACE2 and TMPRSS2 (a host protease required for viral entry) in lung type II pneumocytes, nasal goblet secretory cells, and ileal absorptive enterocytes; these are currently thought to be the target cells of SARS-CoV-2 leading to COVID-19 [15].

Viral isolation and sequencing efforts throughout the world have permitted the interrogation of the SARS-CoV-2 genome, providing a deeper understanding of the evolution of the virus, its proximal origin and revealing patterns of its global spread [16, 17]. These efforts are largely centered on reverse transcription of the viral RNA genome followed by PCR amplicon and hybrid capture based sequencing using Oxford Nanopore and Illumina sequencers [18]. The sequences generated are largely deposited in data repositories such as the sequence read archive (SRA) and GISAID [19]. The availability of these sequences has allowed for platforms such as Nextstrain [20] to continually update and track viral evolution and spread, and to illuminate the viral genetic diversity among carriers [21–24].

At this inter-host level, epidemiological studies often employ per-patient consensus sequences, which summarize each patient’s virus population into a single sequence and ignore minor variants. However, RNA viruses generally exhibit high mutation rates due to error-prone viral RNA polymerases, which typically leads to the presence of various viral variants within a single host [25]. The resulting intra-host diversity has been shown to affect disease progression [26], tissue tropism [27], transmission risk [28], transmission heterogeneity [29], and treatment outcome [30, 31] in various RNA viruses. Recent findings indicate that the mutation rate of SARS-CoV-2 is about as high as the one observed in the SARS-CoV genome (0.80–2.38 × 10^−3^ nucleotide substitutions per site per year) [32, 33]. Although coronaviruses have evolved a proofreading capability attributed to Nonstructural protein 14 resulting in lower mutation rates than other positive-sense ssRNA viruses [34–36], the study of the intra-host genetic diversity of the novel SARS-CoV-2 virus remains important to gain a deeper understanding of its evolution and transmission dynamics and possible implications on its pathology. Previous studies highlight that intra-host genetic diversity in clinical samples is indeed prevalent, and some genomic regions that are susceptible to alterations in the SARS-CoV-2 virus have been identified [37, 38]. Accounting for intra-host diversity can also improve the resolution of phylogenomic analyses of SARS-CoV-2 [39] and our ability to detect selection pressure [40].

To assess the within-host genetic diversity of different viral variants, deep-coverage sequencing is needed to access low-frequency subclonal mutations. However, calling low-frequency mutations reliably from deep sequencing data remains challenging because of various amplification and sequencing errors. Several computational methods have been proposed for this task [41, 42], and specialised bioinformatics pipelines have been developed to streamline and automate the analysis from raw sequencing reads to consensus sequences and single-nucleotide variants [43, 44]. Here, we employ V-pipe [44], a workflow that includes quality control, read mapping and mutation calling, to robustly identify genetic diversity of viral genomes. We examine the intra-host diversity across a large cohort of 3939 public SARS-CoV-2 samples and, as an independent validation cohort, 749 additional samples which have recently been sequenced to study the spread of SARS-CoV-2 in Switzerland [45]. We analyze the distribution of genetic diversity across the genome, including uncovering highly diverse bases and small regions, and we explore the diversity across patient samples. By integrating technical and epidemiological covariates, we show that within-patient viral genetic diversity increases with age of the infected patient.

## Results

We downloaded sequencing data from the SRA from a total of 5934 SARS-CoV-2 samples, which had undergone deep sequencing with Illumina technology. Samples with low (median below 1000) or highly variable (lower quartile below 100 or upper quartile above 10,000) coverage were removed (Figure S1). The remaining 3940 samples were processed using V-pipe [44] (Figure 1a). For each sample, we obtained the coverage across the genome and all mutated positions, including all variant bases and their frequencies (Figure 1b). A single sample with no detected mutations at all was removed to leave a final public cohort of 3939 SARS-CoV-2 samples. In addition, we analyze a new set of 749 sequences derived from samples collected in Switzerland. These sequences were generated as part of our sequencing effort described in [45]. While [45] uses only the consensus sequences for a phylodynamic analysis, here we directly analyse the entire set of raw deep-sequencing reads to assess intra-host diversity. This uniformly produced set of sequences, which we make public on the SRA, serves as a validation cohort.

**Figure 1:**
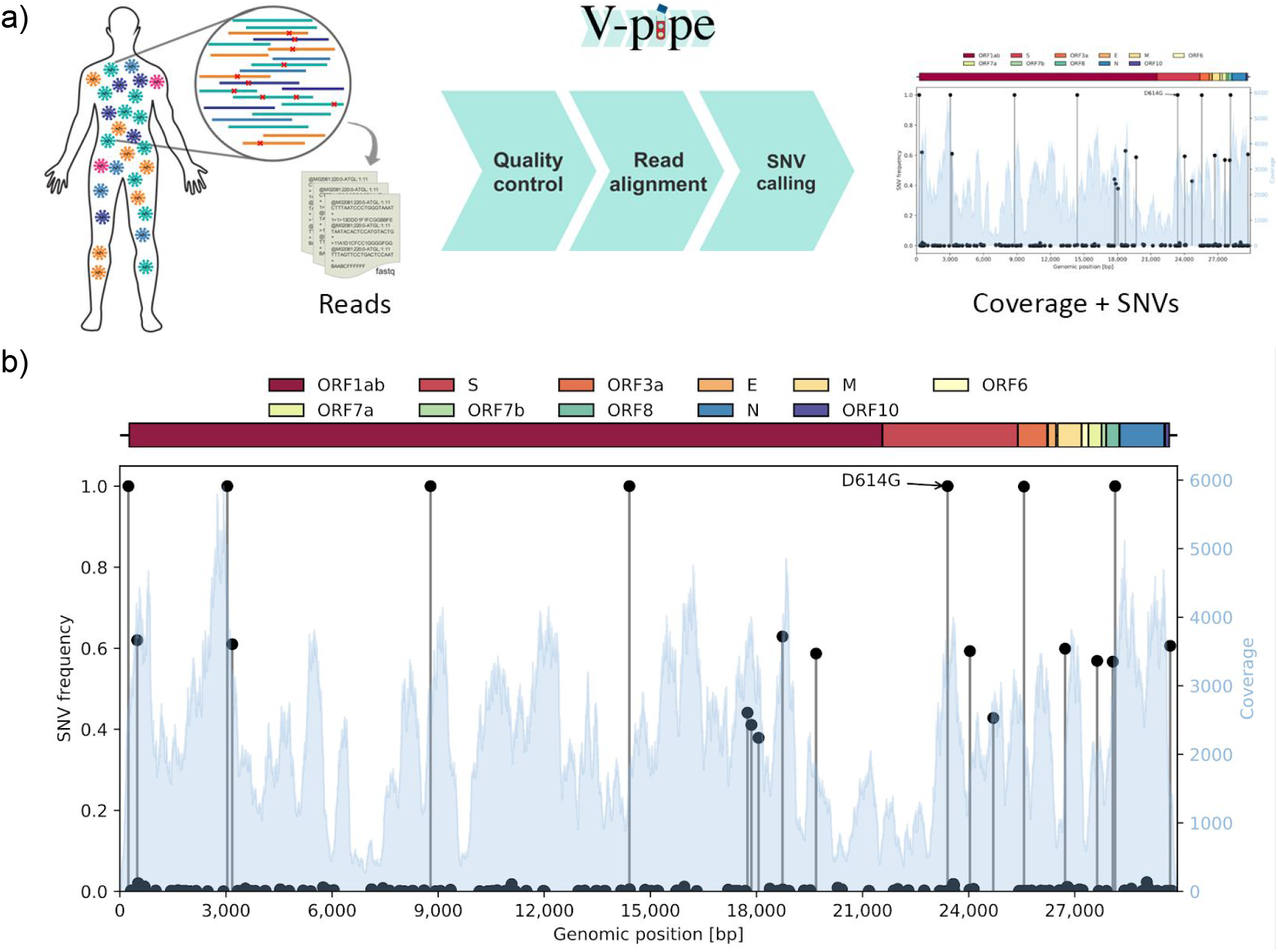
(a) Different viral variants may coexist in the same host such that the sequencing reads contain a mixture of the different components and their SNVs. We employ the bioinformatics pipeline V-pipe for automated end-to-end processing of the raw sequencing data and calling of SNVs and their frequencies [44]. (b) An example output of the workflow for a single sample (SRR11953858) displaying variations in the coverage (blue histogram, right y-axis) and the frequencies of the different SNVs (black lollipops, left y-axis) across the genome (x-axis). Along with clonal mutations with frequencies near 100%, and many low-frequency variants, several other mutations have frequencies of around 40% and 60% in this example.

As a measure of intra-host genetic diversity, we computed the entropy of the nucleotide distribution for each sample and each genomic locus (Materials and Methods). Low entropy indicates a highly conserved site in the patient’s virus population, while high entropy is indicative of variation among nucleotides.

The quartiles of the coverage over the public samples (Figure S2) display typical coverages of over a thousand reads at each genomic site (in line with the coverage filters) meaning that we can detect low-frequency SNVs down to 1-2% [46] (or using estimates from the Lander-Waterman model [47]). However, there are regular dips at the primer locations [48], and noticeable wider dips between genomic positions around 19 to 23 kb. This region covers the end of gene *ORF1ab* encoding the endoribonuclease and 2’-O-methyltransferase, as well as much of the *S* gene producing the S1 subunit, needed for the initial attachment of the virus to the target cell.

We observe a wide spectrum of diversity both across the genome and across samples (Figure 2, focussing on the positions across the genome mutated in at least 4% of the samples). In particular there are subclonal regions visible (as darker contiguous bars in Figure 2) at the start of *ORF1ab*, in the Spike *S* gene and across much of the Matrix *M* gene. There are also highly diverse individual bases and small regions (Tables S1-S4), particularly in the public data.

**Figure 2:**
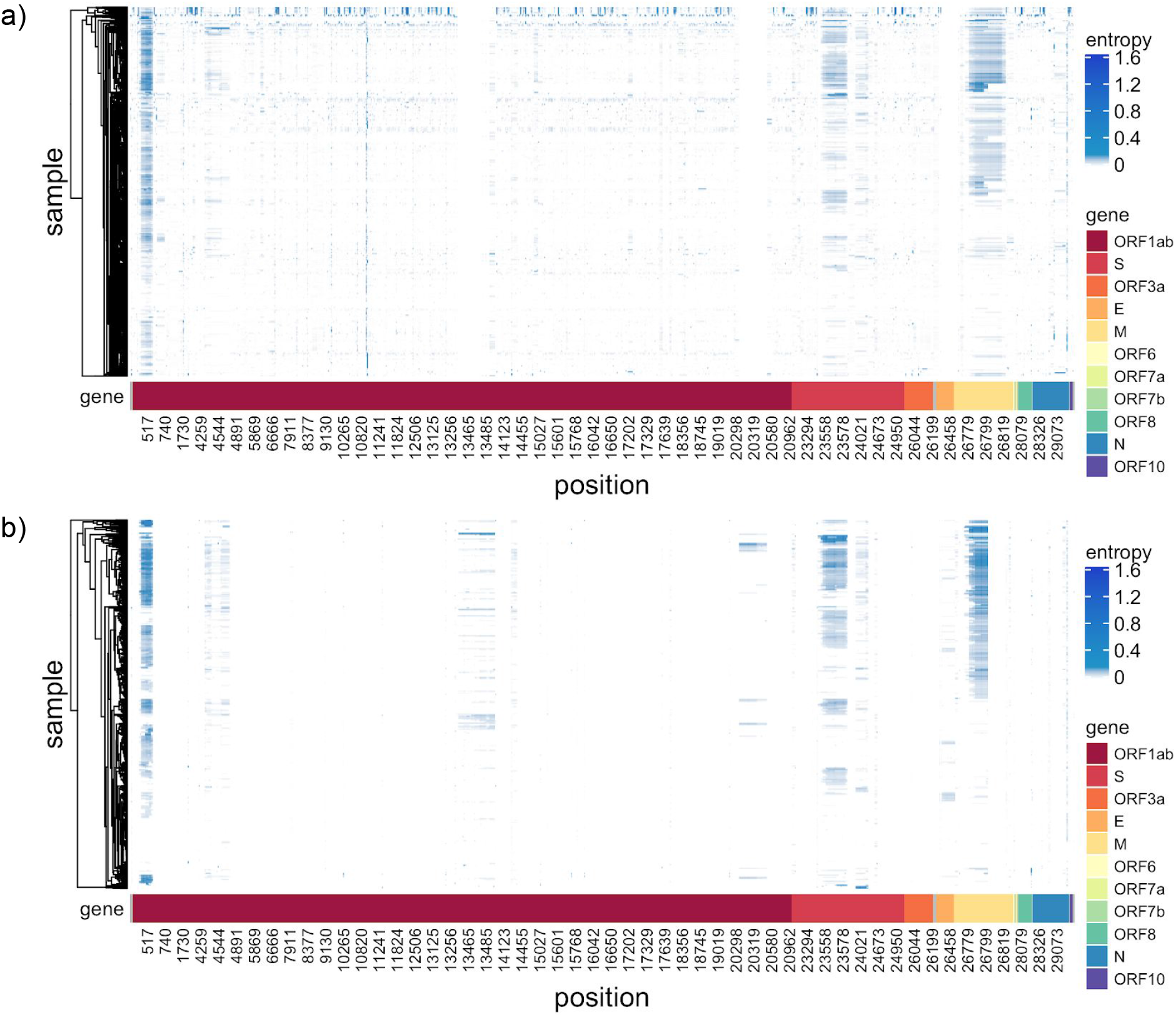
Intra-host diversity measured as entropy of the nucleotide distribution across genome and samples. Only positions with diversity in at least 4% of the samples are selected for each cohort, with the union of positions displayed for comparison. (a) the public cohort, (b) the data from Switzerland.

Next, we summarize genetic diversity per gene by computing the average entropy across all positions of the gene. We observe roughly similar diversity for each gene in the public cohort, except for gene *M* which is much more diverse, and *ORF7b* displays hardly any diversity (Figure 3a), while we observe the same trends, but larger differences across genes in the Swiss cohort (Figure 3b).

**Figure 3:**
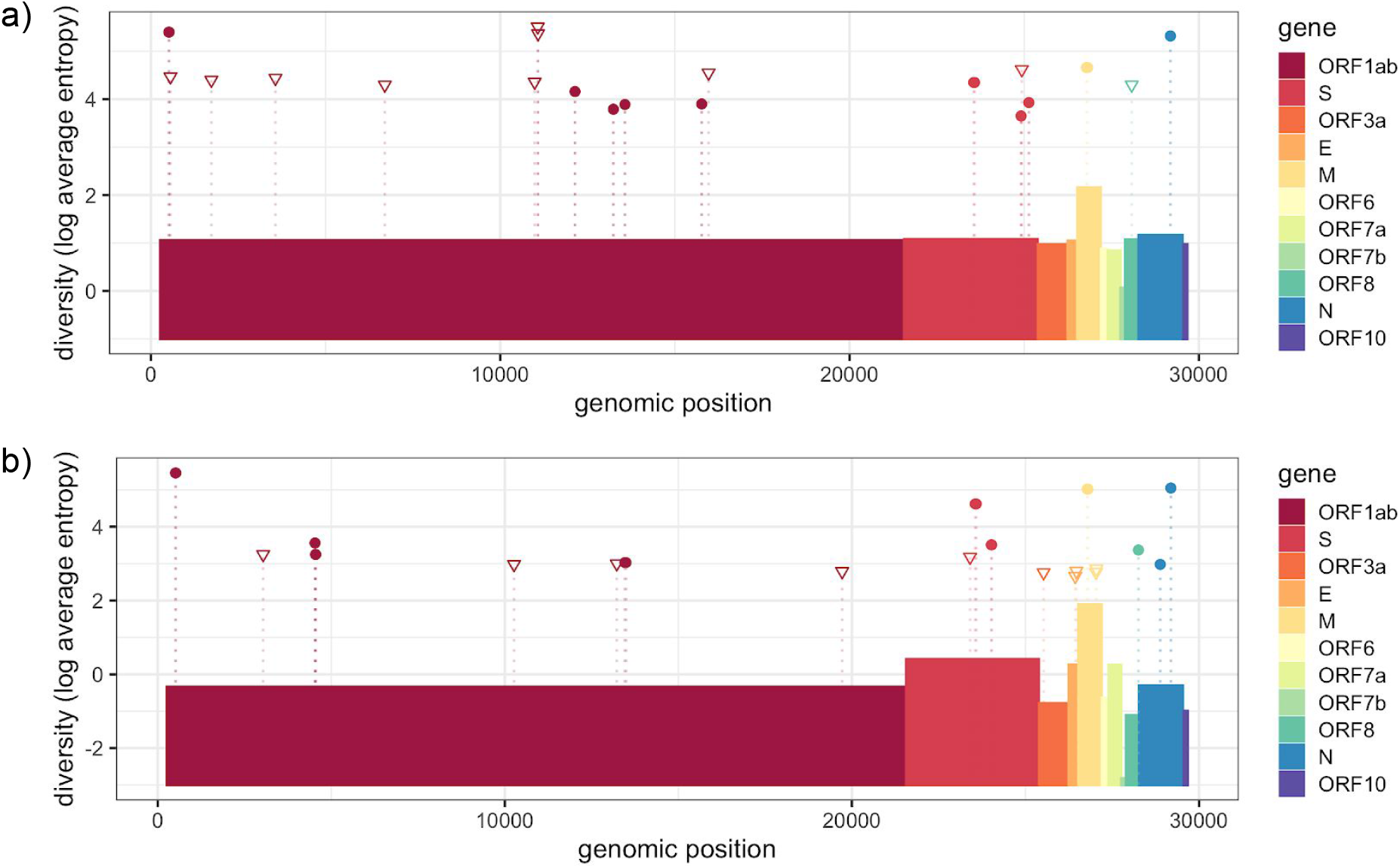
The average entropy per gene (boxes), log transformed, along with the 10 most diverse (in terms of cumulative entropy over samples) positions (triangles) and the 10 most diverse consecutive regions (dots) from Tables S1 and S3 for the public data in (a) and Tables S2 and S4 for the data from Switzerland in (b).

The most diverse site in the public data (Table S1 and Figure 3a), which has clonal or subclonal mutations in nearly half the samples, is at position 11075 in the region of the *ORF1ab* gene coding for non-structural protein 6, a transmembrane protein containing 7 transmembrane helices. Most of the diversity at this position is derived from low-frequency deletions, which would alter downstream amino acids and introduce a premature stop codon. Among the mutated samples, deletions occur at an average rate of 2.62%. 12.5% of mutated samples include variants with a T>C substitution, which occurs with an average frequency of 1.75% among those samples (corresponding to an overall average rate of 0.22% among all mutated samples). In the reference genome, the codon incorporates a Phenylalanine amino acid (Phe35 of nsp6), however, any variation here will lead to a change in the inserted amino acid; for example, a T>C mutation results in the incorporation of a Leucine.

Another residue in close proximity to Phe35, namely Leu37 of nsp6, has often been found to be replaced by a Phenylalanine residue in recent sequences from Europe, Asia and America [49]. Here, we also identify a high entropy at position 11083 inside the codon of Leu37 (Table S1), which is affected in around a quarter of samples. Any alteration from the reference base G to either a T or C results in a Phenylalanine residue as opposed to a Leucine. Here we observe a large average frequency of Thymine bases (44.65%) and common deletions (4.93%) which also lead to a Phenylalanine residue and a premature stop codon.

Genomic position 23403 is the site at which an A>G mutation causes the well-known D614G amino acid alteration in the Spike region. Studies suggest that this alteration provides a fitness advantage and becomes the more prevalent variant in the population over time [50], and it is also the dominant variant in our public cohort. We find that it is among the 100 most diverse positions in the SARS-CoV-2 genome (rank 95 in our cohort) with 36.0% of our samples having mutations there relative to our cohort consensus G, and 29.7% having diversity of different variants coexisting. Among the mutations, there is an average frequency of 90.4% A, 9.5% G and 0.1% deletions. In the Swiss cohort, position 23403 has the second highest diversity among all individual bases (Table S2).

We also assessed genetic diversity over consecutive genomic regions (Materials and Methods). The most diverse regions in both the public and Swiss data (Tables S3 and S4, adjacent sites all with a log entropy above 3.0 for the public data and above 2.0 for the Swiss data) include a span of 16 bases at the start of the *ORF1ab* gene (Figure 3). This region forms part of the nsp1 protein which shuts down host mRNA translation and protein production via interactions with the 40S and 80S ribosomal subunits. A notable consequence of this nsp1 induced inhibition of translation and protein production is the reduction in innate immune response activity, specifically interferon signalling [51].

The next most diverse genomic region (Tables S3 and S4) consists of two adjacent positions at 29187 and 29188 within a single codon in the *N* gene forming the nucleoprotein. The reference bases at those two positions are C and A, respectively, which results in the incorporation of an Alanine residue at position 305 of the nucleoprotein (Ala305). However, due to the redundancy of the genetic code, alterations in position 29188 alone do not lead to an alteration on the protein level.

The remaining regions in Table S3 with a length greater than two lie within the *M* and *S* genes, and are similarly present in the Swiss cohort (Table S4). The 41 nucleotides from 26780 to 26820 lead to the incorporation of 15 amino acids from Cys86 to Phe100 forming a section of the matrix protein’s transmembrane region. The matrix protein is known to be central in viral assembly with several interacting partners including itself, the envelope protein, the nucleoprotein and the spike protein [52, 53]. The spike protein, when primed, is divided into the S1 and S2 subunits with S1 being essential for hACE2 recognition and S2 mediating viral entry into the host cell. Another region of high entropy comprises 30 nucleotides within the Spike gene which incorporates 11 amino acids to the S1 subunit of SARS-Cov-2 ranging from Ile664 to Tyr674. These amino acids lie between the receptor binding domain and Furin cleavage site needed for S protein priming [54, 55].

The distribution of total entropy per base has a long tail (Figure 4a) resulting in locations with extremely high diversity, such as the examples discussed above, while the majority of the genome is relatively conserved. Taking a logarithmic transform of the entropy we can observe much more detail in the distribution, with a roughly normal distribution and a slight negative skew in the public data (Figure 4a inset). Considering the entropy per sample instead provides a picture of how diversity is distributed among samples. We again find a long tail (Figure 4b) with certain individuals having vastly more internal diversity than the majority of samples dominated by clonal variants and low-frequency mutations. Detecting diversity in each sample is heavily dependent on the sequencing technology, sample processing, and sequencing depth. For example, all the most diverse samples (Table S5) come from one SRA study (SRP253798) which makes up 29% of the entire cohort. The samples with the highest detected diversity had between 10% and 30% of the genome affected, with the vast majority of their mutations (over 99.8%) being subclonal and the coexistence of more than one character (base or deletion) in the mapped reads.

**Figure 4:**
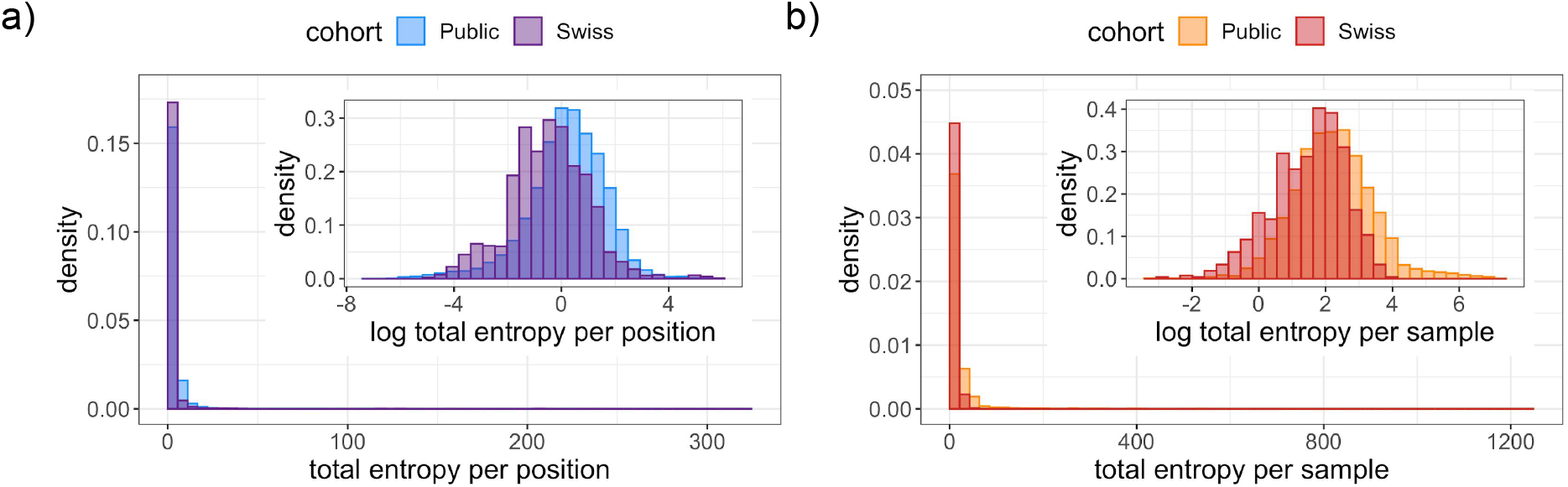
The distributions of the total entropy in the public and Swiss cohorts per position (a) and per sample (b). Insets: Under a logarithmic transformation, the distribution per position (a) and per sample (b).

Like the entropy, the number of clonally or subclonally mutated positions per sample also has a long tail (Figure S3) with, in the public data, for example, quartiles at 119, 211 and 378.5 positions being mutated (corresponding to 0.40%, 0.71% and 1.27% of the genome) but a maximum of 8705 positions (29.11% of the genome).

Finally, we tested the hypothesis that epidemiological parameters are related to viral genetic diversity. Specifically, to determine whether the host’s age, sex or geographical location predict the diversity of their virus population, we performed regression modelling on the subset of samples for which we have such information. This resulted in 1043 samples from Australia which were sequenced with paired-end amplicon sequencing with PCR amplification. We also adjusted for technical parameters of the sequencing to avoid confounding (Materials and Methods). We found that sex is not a significant predictor of diversity (*p* = 0.50). By contrast, age is significantly associated with intra-host viral genetic diversity (*p* = 4 × 10^−4^) with each decade increasing the total entropy by 8.6% on average (Table 1, Figure S5).

**Table 1:** regression coefficients of log entropy against the above predictors for the 1043 public samples with covariate information.

| Predictor | Coefficient | Standard error | t-statistic | p-value |
| --- | --- | --- | --- | --- |
| sex (male) | -0.0563 | 0.0839 | -0.671 | 0.50 |
| age (decade) | 0.0822 | 0.0231 | 3.56 | 0.00038 |
| log(median coverage) | 0.190 | 0.125 | 1.51 | 0.13 |
| log(IQR/m) | 1.48 | 0.122 | 12.1 | $< 2 \cdot 10^{-16}$ |

Clinical covariates were not available for the Swiss cohort, but sequencing date was utilized as a proxy for decreasing age, since testing expanded to younger populations over time (Figure S6). The significant decrease in diversity over time (*p* < 10^−7^; Table 2, Figure S7) in the Swiss cohort therefore corroborates the association of increased diversity with age uncovered in the public cohort.

**Table 2:**
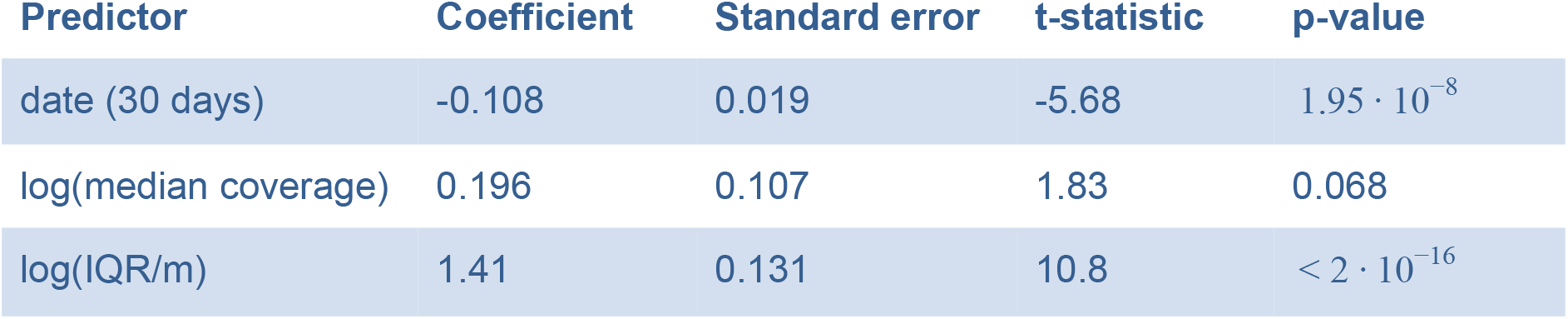
regression coefficients of log entropy against the above predictors for the 749 Swiss samples.

## Discussion

We processed a large cohort of 3939 deeply sequenced public SARS-CoV-2 genomes, and 749 samples from Switzerland, with the bioinformatics software V-pipe [44] to uncover within-patient genetic diversity. We observe a heavy tail distribution in diversity per sample and per position indicating that much of the diversity is concentrated in small numbers of sites and patients. The most diverse small regions were consistent between the public and Swiss data, though the individual bases were more varied.

Detecting and quantifying intra-patient genetic diversity from deep sequencing data is technically challenging. It may be heavily influenced by the sample preparation and how it is sequenced, along with possible artifacts arising from the process, so that extremely diverse patients may not be comparable across cohorts. Accounting for such technical parameters of the sequencing process and coverage, which affects the detection limit of diversity, we find that age is a significant predictor of diversity in the public data. The model predicts that on average genetic diversity increases by 8.6% every ten years. The increase in diversity with host age is corroborated in the Swiss data. Age has previously been associated strongly with worse disease outcome and higher death rates [56], along with concomitant comorbidities [57]. With high-quality clinical and genetic data, it will be highly relevant to see whether diversity is a cause or a consequence of disease progression. Likewise, with future transmission network data it will be interesting to uncover whether diversity increases infectiousness, as for influenza [28].

The detection of subclonal mutations is affected by the sequencing depth at each position, and across the genome depending on the amplification and capture of RNA for sequencing some regions may be more poorly resolved. The coverage distribution therefore affects our ability to detect highly diverse bases. With this caveat, the most diverse gene is the Matrix *M* gene while highly diverse positions include a mix of low-frequency variants common to a quarter of the cohort or more, and rarer high-frequency subclonal mutations in around 5% of the cohort. The observation of common low-frequency and less common high-frequency genetic variants is in line with previous research on both intra- and inter-host genetic diversity of SARS-CoV-2 [21, 39]. The D614G variant, which appears to increase infectivity and is becoming more dominant over time [50] is the dominant variant in our public cohort. It also exhibits high intra-host diversity with 29.7% of the cohort experiencing subclonal mutations with the different variants coexisting. This diversity is mimicked in the data from Switzerland, where the D614G variant is actually encoded by the second most diverse genomic position.

## Materials and Methods

### Public data

We retrieved data from the Sequence Read Archive (https://www.ncbi.nlm.nih.gov/sra) on June 10, 2020. Samples with the term ‘“Severe acute respiratory syndrome coronavirus 2”[Organism] OR “Sars-Cov-2[All Fields]”’ were filtered and only one copy of duplicates with the same BioSample ID was retained. We used the meta file to further filter the samples by Illumina technology. This resulted in 5934 samples which were associated with downloadable data.

Subsequent to downloading the selected sample set, we trimmed all read files using PRINSEQ ([58] version 0.20.4, parameters: -ns_max_n 4 -min_qual_mean 30 -trim_qual_left 30 -trim_qual_right 30 -trim_qual_window 10 -min_len <80% of average read length>), mapped them to NC_045512.2 using bwa ([59] version 0.7.17-r1188, subcommand: mem). Coverage quartiles for each sample are displayed in Figure S1.

### Data from samples collected in Switzerland

Sample collection and sequencing are detailed in [45]. Briefly, we obtained RNA samples extracted from nasal swab tests which had previously tested positive on RT-qPCR from Viollier AG laboratory and sequenced them at the Genomics Facility Basel. We performed reverse transcription using random hexamers and PCR amplified the resulting DNA with primers from the artic-ncov2019 protocol [https://github.com/artic-network/artic-ncov2019/tree/master/primer_schemes/nCoV-2019/V3]. We prepared libraries from these 4000 bp-long amplicons using Illumina TruSeq adapter sequences and sequenced them on an Illumina MiSeq System (Paired-end sequencing, 2x 251 cycle). The phylogenetic relationship between the consensus sequence of 681 samples from this collection has been previously analyzed as part of the subset of GISAID data available for Switzerland until July 10, 2020 in [24] and results based on the consensus sequences of the full dataset are presented in [45]. Here we focus on the raw reads directly and analyse the deep sequencing data to uncover within host diversity. The collection of swabs analysed here spans a time period from Mar 4, 2020 to August 13, 2020, with samples from across Switzerland. The analysed reads are available in the SRA (see Data Availability below).

Data on the age distribution of COVID-19 cases in Switzerland was downloaded from [https://www.bag.admin.ch/dam/bag/de/dokumente/mt/k-und-i/aktuelle-ausbrueche-pandemien/2019-nCoV/covid-19-basisdaten-fallzahlen.xlsx.download.xlsx/Dashboards_1&2_COVID19_swiss_data_pv.xlsx] on September 21, 2020.

### Filtering

Samples were subset by applying a coverage filter (minimum lower quartile: 100, minimum median: 1000, maximum upper quartile: 10000). After this filtering, 3940 public samples and 749 samples from Switzerland were retained for later analyses.

### Data processing

We used V-pipe ([44]; sars-cov2 branch of https://github.com/cbg-ethz/V-pipe) to call variants for each sample using ShoRAH [60] and default settings, including discarding deletions with a frequency below 0.5%. One sample had no remaining variants detected and was excluded giving a final cohort size of 3939. For each called variant per sample, we computed the relative frequency *f_k_* of each character *k* (nucleotide A, C, G, T, or deletion) among the mapped reads at that position. The entropy is then computed as

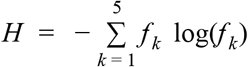

summarising the five frequencies in a single measure of diversity. The entropy is zero whenever one character has a frequency of 100%, and is maximised when all the characters have equal frequency 1/5, giving a maximum value of log(5) ≈ 1.61. We denote the entropy of sample *i* at base *j* by *H_ij_* and compute the total entropy per position in the genome by summing over samples 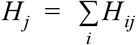 (Table S1 and S2). For the data from Switzerland we additionally multiply by the ratio of cohort sizes (3939/749) to make the total entropy values per position comparable across the cohorts. We compute the average entropy over a consecutive genomic region *J* as 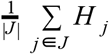, and take the logarithm to obtain the log average entropy (Table S3 and S4). We compute the total entropy per sample by summing over bases 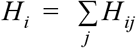 (Table S5).

### Regression modelling

To evaluate which covariates are predictive of diversity, we build a regression model on the public data of log total entropy on age, sex, and country of origin. To adjust for the possible effects of coverage and sequencing technology on the diversity, we include factors for paired-end or single-end sequencing, the assay type, the library selection and the SRA study. We also include the logarithm of the median coverage. Since not only the average coverage, but its variability may affect the ability to detect SNVs, we further include the IQR of the coverage, divided by the median *m*, and again log transformed:

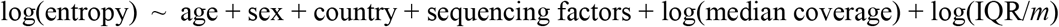

Filtering the samples which have age and sex information left 1060 samples, of which all but 17 were from Australia. We therefore retained just those 1043 from Australia, and removed the country dependence from the regression. All remaining samples were paired-end amplicon sequencing with PCR amplification from the study SRP253798, so those factors were also removed to provide the final regression:

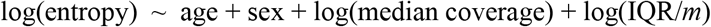

Of the 1043 samples, 480 (46.0%) were female and 563 (54.0%) were male, while the distribution of ages (Figure S4) has a median value of 46 and lower and upper quartiles at 29 and 60. The results of the regression are listed in Table 1.

For the data from Switzerland, we do not have clinical covariates, only the date of sequencing. However, we can use the sequencing date as a proxy for age, because over time, the age distribution has progressively reduced in Switzerland as a whole (Figure S6). All samples were processed in the same way, so for the regression we model

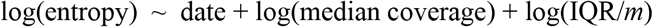

with the results in Table 2.

For visualisation purposes, we regress patient age on sample date from data collected about positive tests in Switzerland as a whole (Figure S6). The model is then used to predict the age of our 749 deeply sequenced samples based on the sequencing date. Against the predicted age, we plot the log total entropy, adjusted for the coverage covariates, to show how the negative correlation with date (Table 2) corroborates an increase in entropy with age (Figure S7).

## Code availability

The source code to process the samples and perform and reproduce the analyses is available on GitHub (https://github.com/cbg-ethz/SARS-CoV-2_Analysis) in the form of multiple Snakemake [61] workflows.

## Data availability

The public data is available from the SRA, as described in the Methods and Materials section. The Swiss data has been added to the SRA with accession number PRJEB38472, scheduled to be publicly available from November 1, 2020.

## Author contributions

Conceptualization: JK, NB; Data Curation: KPJ, MP, IT, SN, SS, NSdS; Formal Analysis: JK, KPJ, KJ; Funding Acquisition: TS; Methodology: JK, KPJ, NB; Resources: IN, NS, EB, TSc, VC, CBec, OK, CN, MR, CB; Software: JK, KPJ, AB, NBo, FB, AD, PF, KJ, LL, MP, SPC, IT; Supervision: NB, TS; Visualization: JK, AB, M-AD, KPJ; Writing – Original Draft Preparation: JK, AB, KPJ, FB; Writing – Review & Editing: all authors.

## Supplementary Figures and Tables

**Figure S1.**
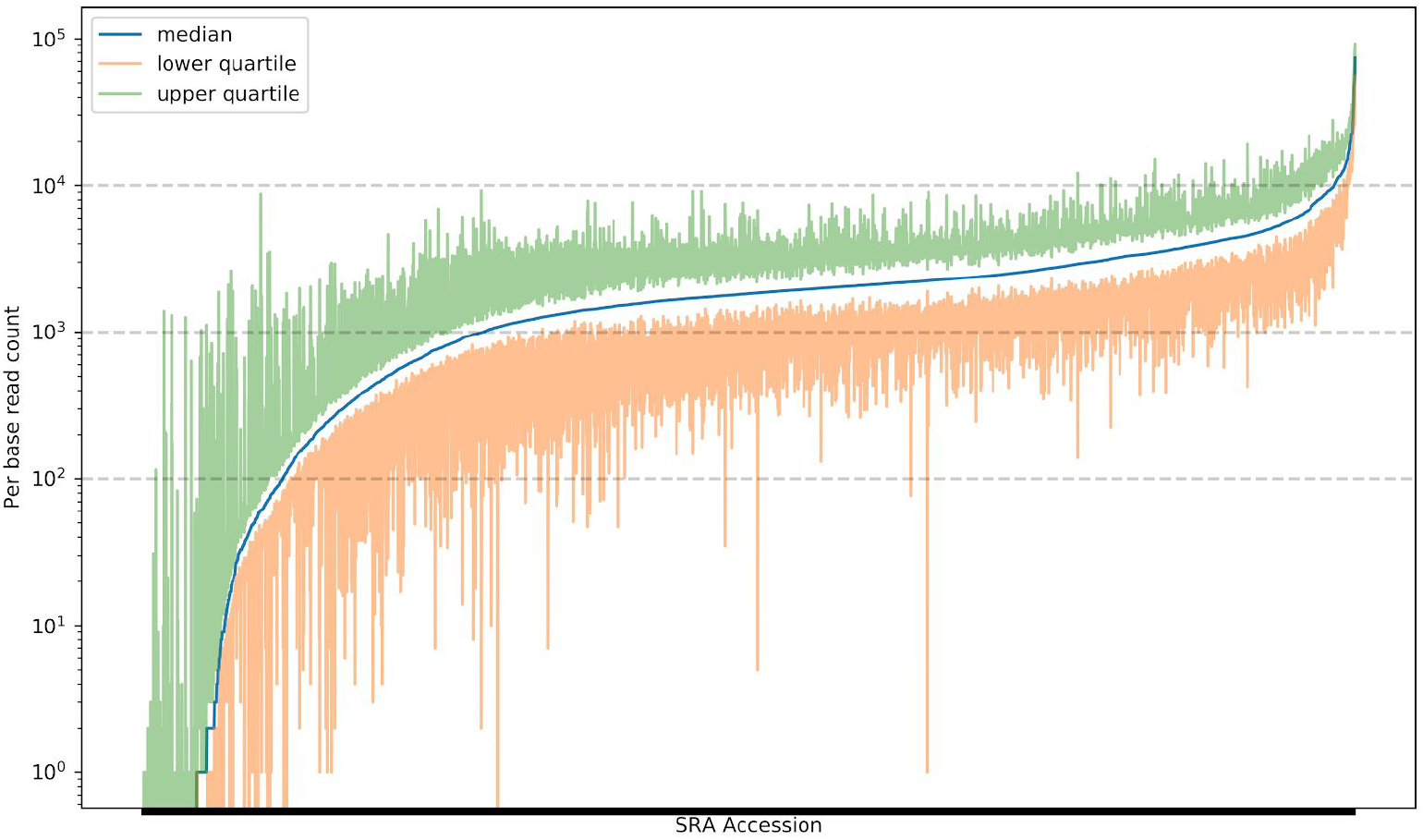
The median coverage, along with the lower and upper quartiles for the 5934 samples downloaded from the Sequence Read Archive. The dashed grey lines correspond to the coverage filters used to subset the samples before further processing (minimum lower quartile: 100, minimum median: 1000, maximum upper quartile: 10000).

**Figure S2:**
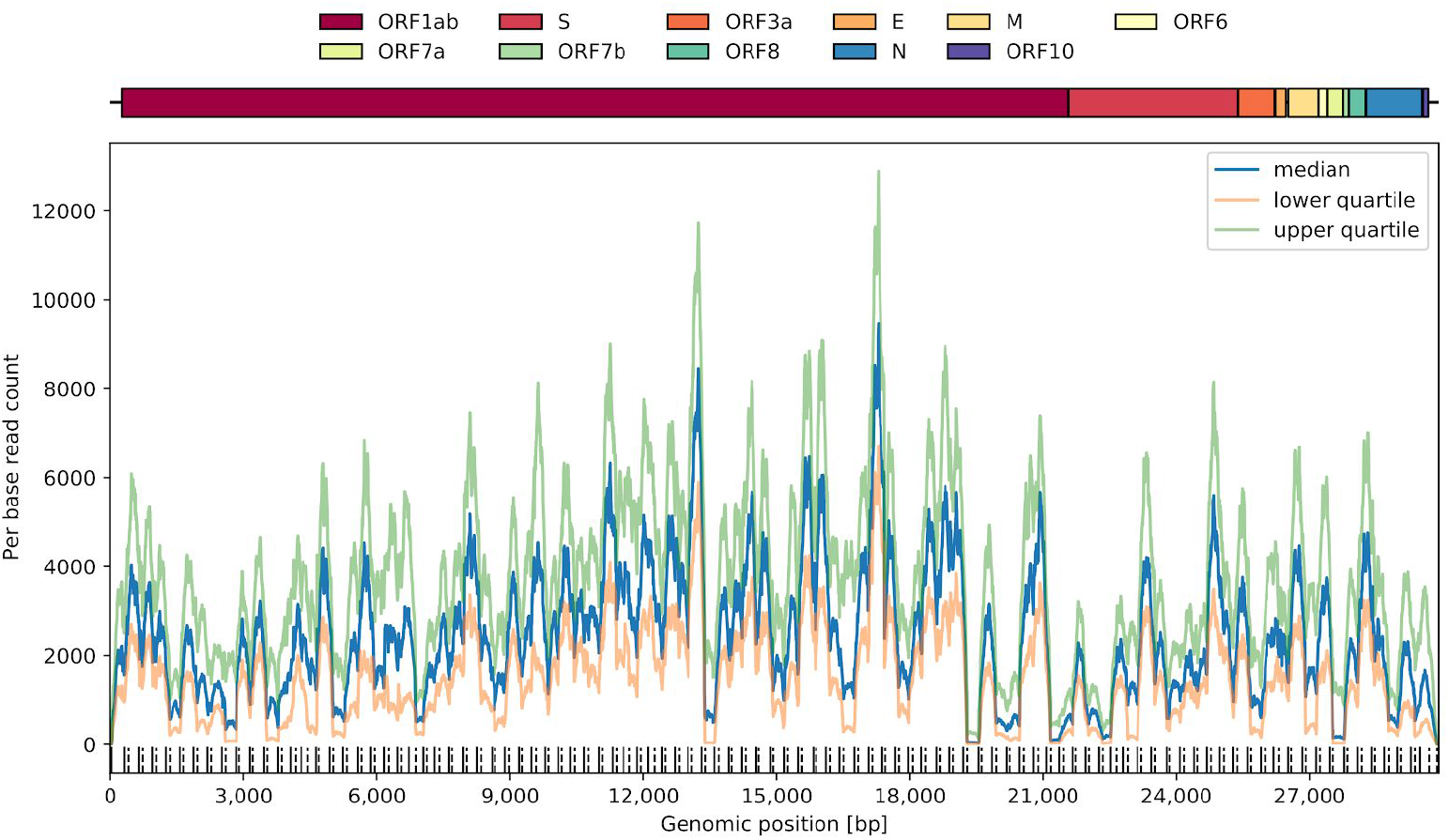
The lower and upper quartiles of the coverage per genomic position, along with the median coverage, among the 3940 coverage-filtered public sample sequences. The locations of typical primers are indicated along the bottom row (solid at the start of left primers and dashed at the end of right primers).

**Table S1:** The 10 most diverse positions in the genome in the public data ranked by their entropy, along with the fraction of samples exhibiting any mutation, and, for those samples, the distribution of average mutation frequencies across the different bases or deletion.

| position | gene | log<br>entropy | mutated<br>samples (%) | ref.<br>base | A<br>(%) | C<br>(%) | G<br>(%) | T<br>(%) | del.<br>(%) |
| --- | --- | --- | --- | --- | --- | --- | --- | --- | --- |
| 11075 | ORF1ab | 5.51 | 49.30 | T | 0.00 | 0.22 | 0.00 | 97.16 | 2.62 |
| 11083 | ORF1ab | 5.37 | 24.37 | G | 0.01 | 0.01 | 50.40 | 44.65 | 4.93 |
| 24933 | S | 4.62 | 6.93 | G | 0.02 | 0.92 | 78.08 | 20.54 | 0.44 |
| 15965 | ORF1ab | 4.55 | 28.21 | G | 0.00 | 0.00 | 98.31 | 1.35 | 0.34 |
| 558 | ORF1ab | 4.47 | 5.97 | G | 0.04 | 0.41 | 82.82 | 16.37 | 0.36 |
| 3564 | ORF1ab | 4.44 | 4.32 | G | 0.08 | 0.34 | 64.54 | 34.63 | 0.42 |
| 1730 | ORF1ab | 4.40 | 24.45 | G | 1.84 | 0.00 | 98.16 | 0.00 | 0.00 |
| 10986 | ORF1ab | 4.36 | 5.69 | G | 0.91 | 0.27 | 88.20 | 9.19 | 1.43 |
| 6696 | ORF1ab | 4.30 | 22.34 | C | 0.01 | 96.92 | 0.00 | 2.97 | 0.10 |
| 28079 | ORF8 | 4.30 | 4.72 | G | 0.00 | 0.13 | 83.48 | 14.62 | 1.76 |

**Table S2:**
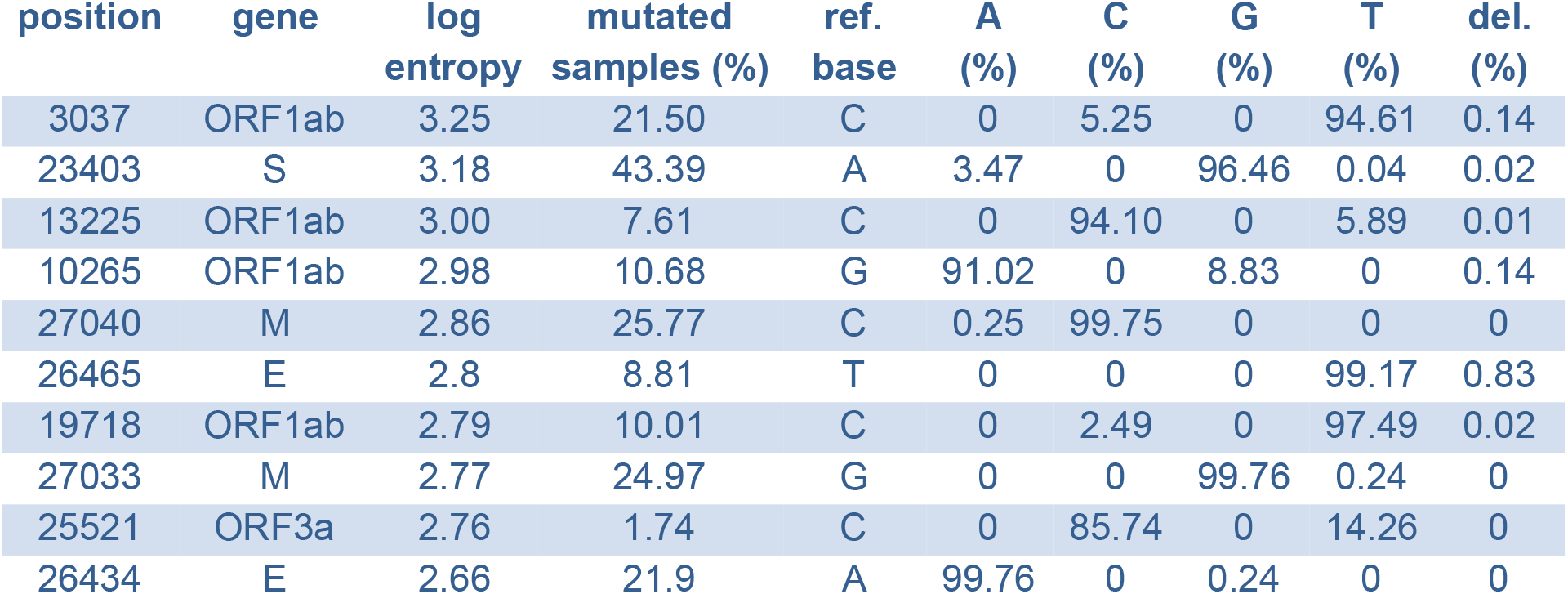
The 10 most diverse positions in the genome in the Swiss data ranked by their entropy, along mutation patterns for those positions across the cohort.

| position | gene | log<br>entropy | mutated<br>samples (%) | ref.<br>base | A<br>(%) | C<br>(%) | G<br>(%) | T<br>(%) | del.<br>(%) |
| --- | --- | --- | --- | --- | --- | --- | --- | --- | --- |
| 3037 | ORF1ab | 3.25 | 21.50 | C | 0 | 5.25 | 0 | 94.61 | 0.14 |
| 23403 | S | 3.18 | 43.39 | A | 3.47 | 0 | 96.46 | 0.04 | 0.02 |
| 13225 | ORF1ab | 3.00 | 7.61 | C | 0 | 94.10 | 0 | 5.89 | 0.01 |
| 10265 | ORF1ab | 2.98 | 10.68 | G | 91.02 | 0 | 8.83 | 0 | 0.14 |
| 27040 | M | 2.86 | 25.77 | C | 0.25 | 99.75 | 0 | 0 | 0 |
| 26465 | E | 2.8 | 8.81 | T | 0 | 0 | 0 | 99.17 | 0.83 |
| 19718 | ORF1ab | 2.79 | 10.01 | C | 0 | 2.49 | 0 | 97.49 | 0.02 |
| 27033 | M | 2.77 | 24.97 | G | 0 | 0 | 99.76 | 0.24 | 0 |
| 25521 | ORF3a | 2.76 | 1.74 | C | 0 | 85.74 | 0 | 14.26 | 0 |
| 26434 | E | 2.66 | 21.9 | A | 99.76 | 0 | 0.24 | 0 | 0 |

**Table S3:**
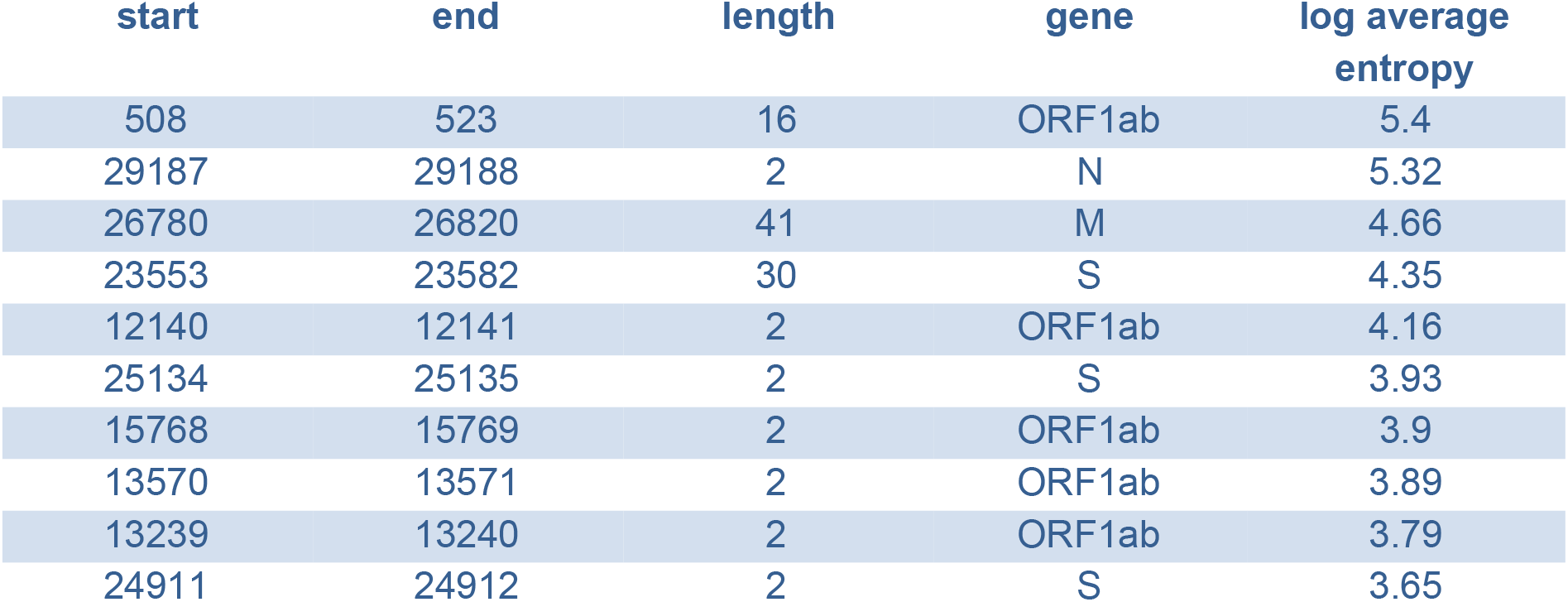
The 10 most diverse consecutive regions in the genome in the public data, ranked by their average entropy.

**Table S4:**
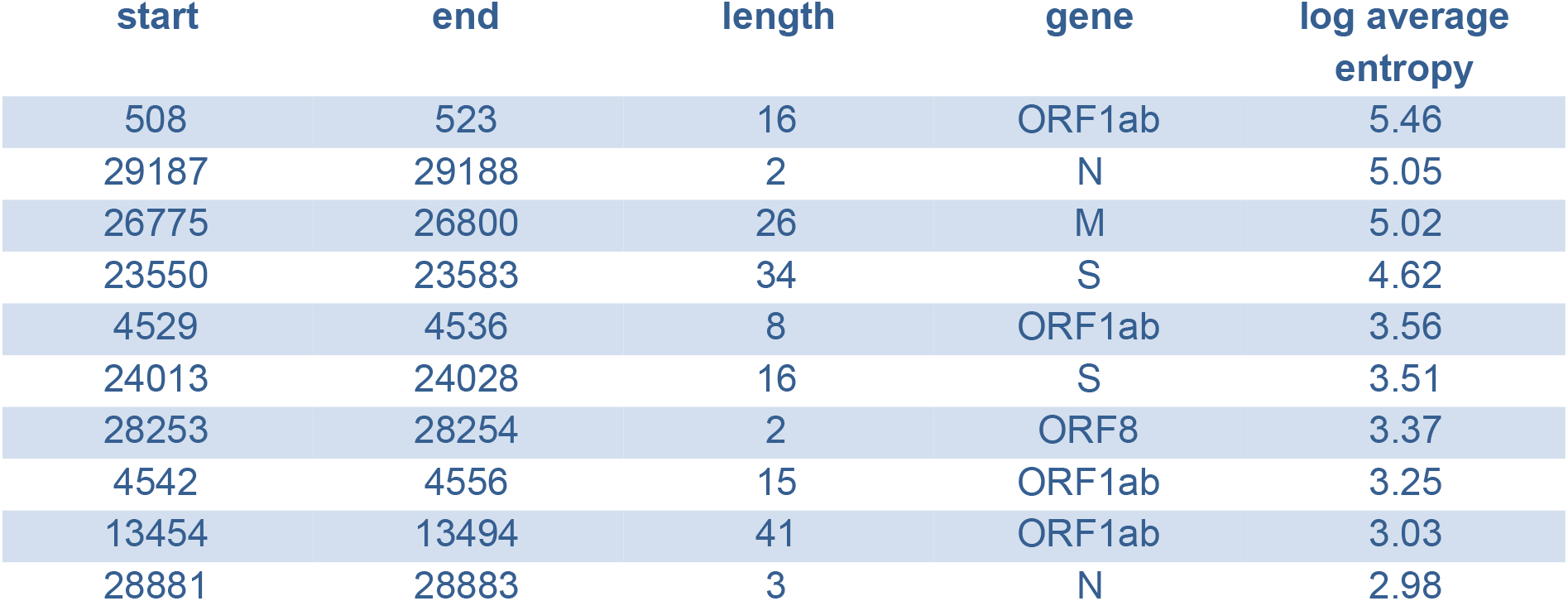
The 10 most diverse consecutive regions in the genome in the Swiss data, ranked by their average entropy.

**Table S5:** The 10 samples with the highest measured diversity, ranked by their entropy, and the fraction of their positions affected by mutations.

| sample ID | log entropy | mutated positions (%) | sample ID | log entropy | mutated positions (%) |
| --- | --- | --- | --- | --- | --- |
| SRR11577862 | 7.08 | 17.41 | SRR11578392 | 6.82 | 18.40 |
| SRR11578151 | 7.03 | 29.11 | SRR11578356 | 6.76 | 19.23 |
| SRR11577865 | 6.97 | 19.92 | SRR11578031 | 6.67 | 11.54 |
| SRR11578029 | 6.90 | 14.60 | SRR11577858 | 6.61 | 17.94 |
| SRR11578028 | 6.88 | 20.05 | SRR11578137 | 6.54 | 19.26 |

**Figure S3:**
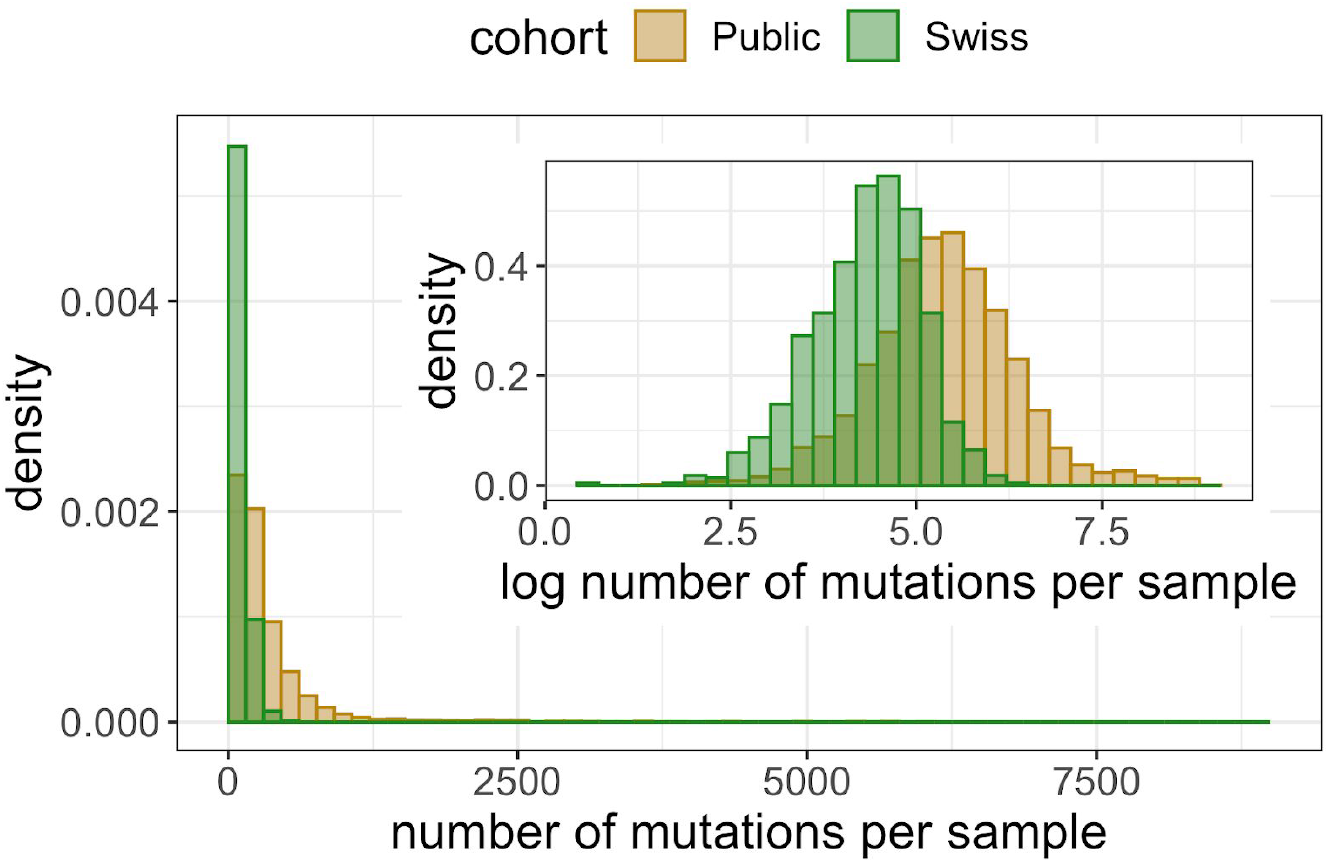
The distributions of the number of mutated positions per sample. Inset: the distribution under a logarithmic transform.

**Figure S4:**
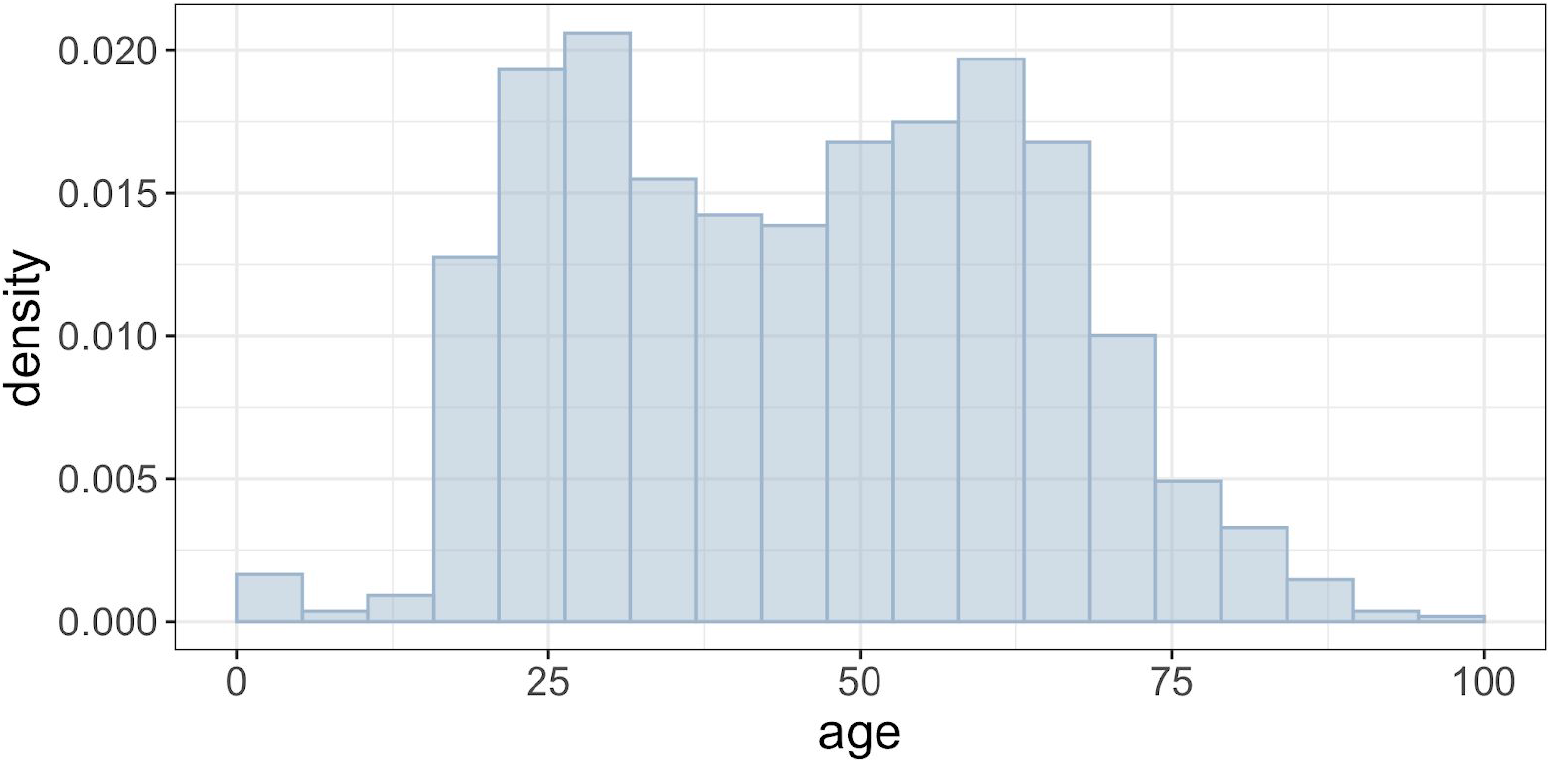
The distributions of ages of the 1043 public samples in the regression modelling.

**Figure S5:**
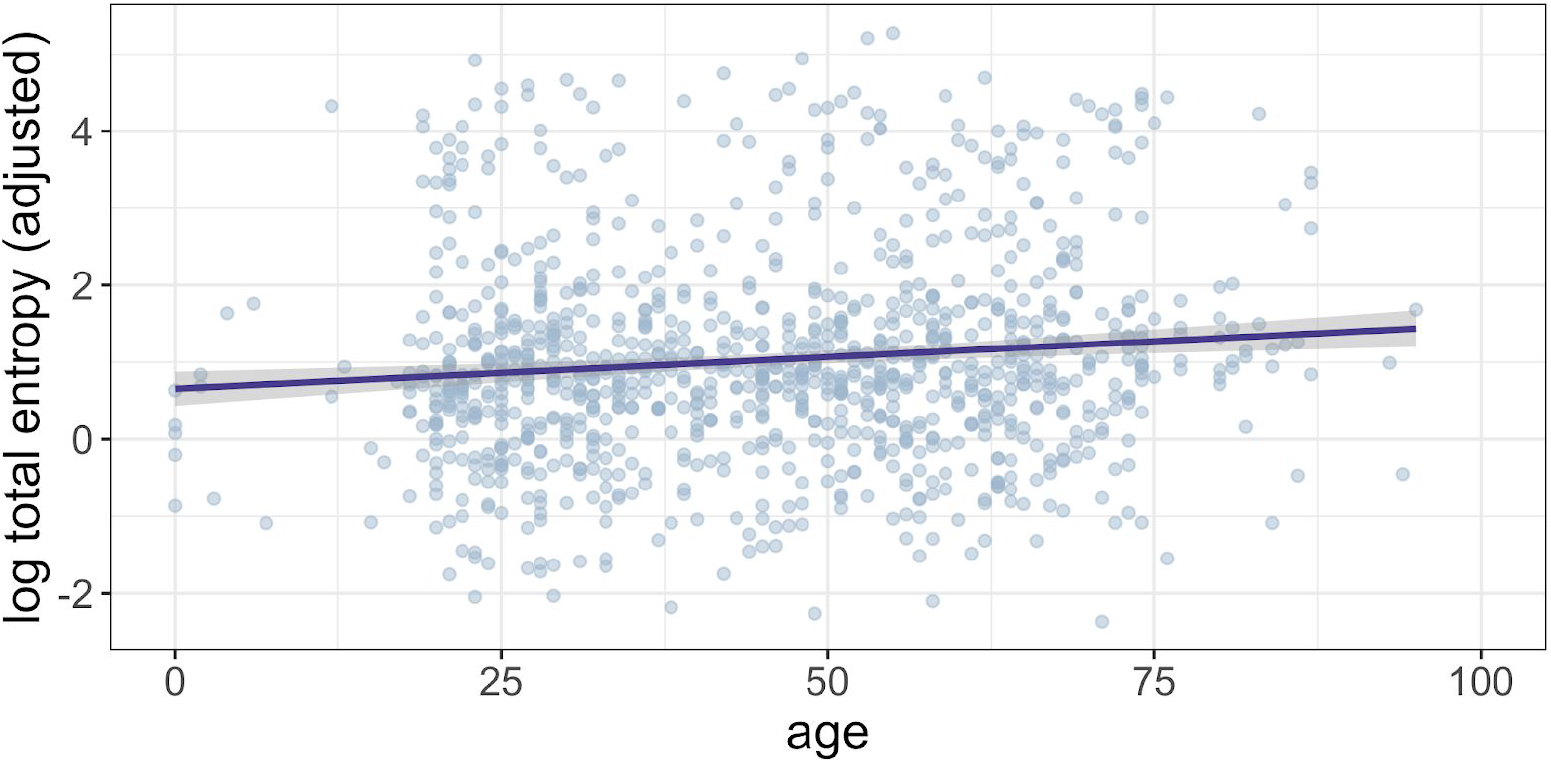
The dependence of log total entropy on age for the 1043 public samples, after adjustment for sex and sequencing coverage covariates in the regression modelling.

**Figure S6:**
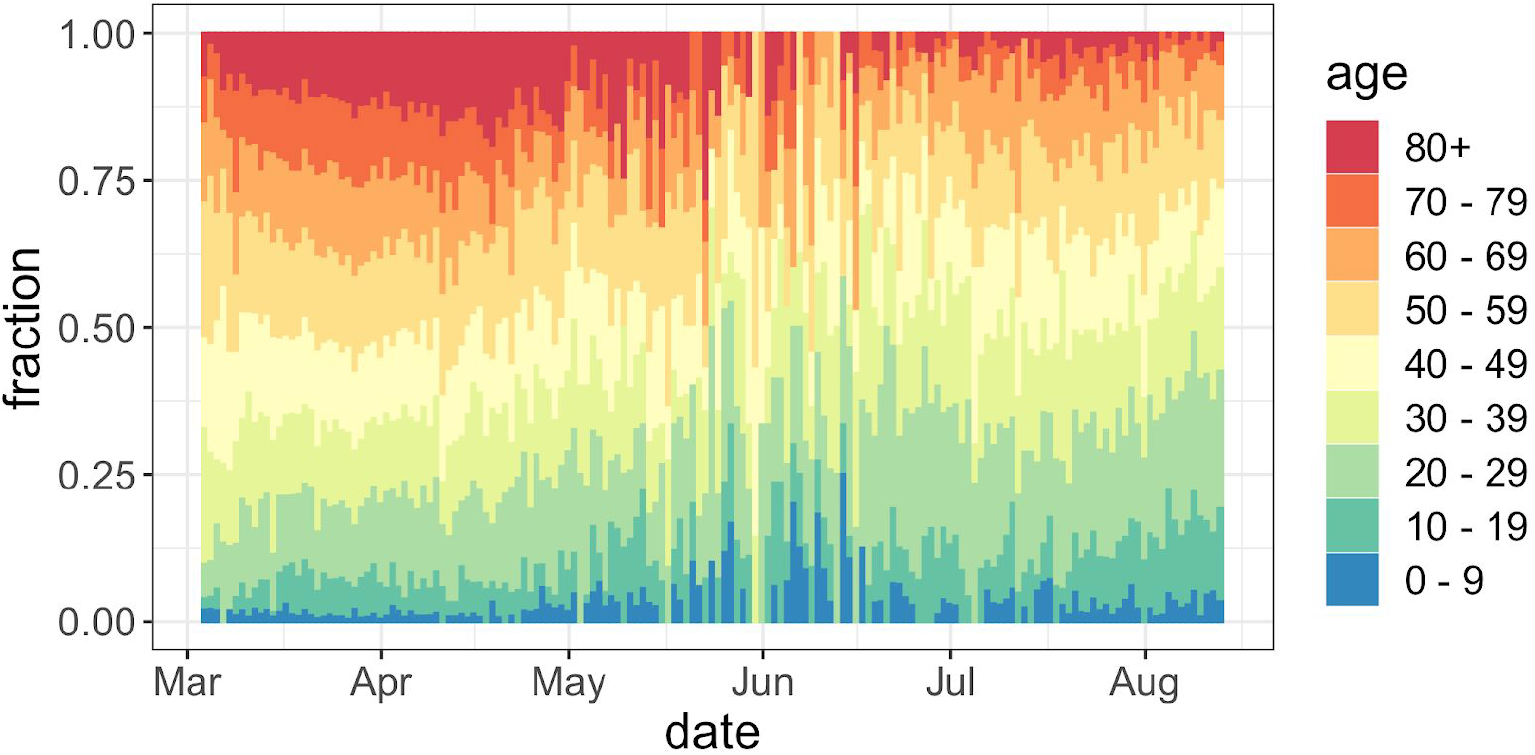
The distribution of ages of positive Covid-19 tests in Switzerland as a whole, covering the period where our cohort of deeply sequenced samples was collected.

**Figure S7:**
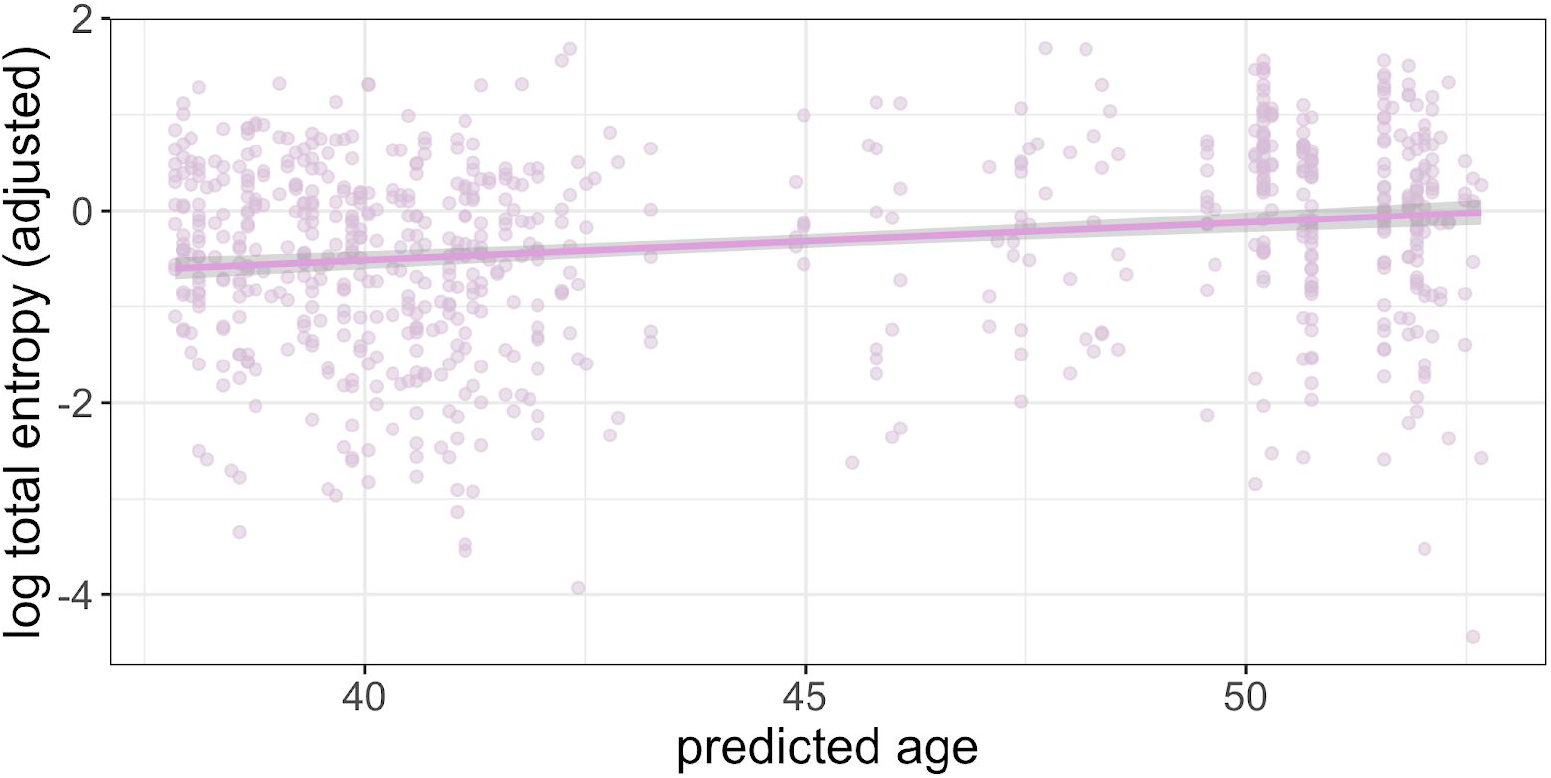
The dependence of log total entropy on predicted age for the 749 Swiss samples, after adjustment for sequencing coverage covariates in the regression modelling. The predicted age is constructed from a linear model of age on date built with the data displayed in Figure S6.

